# An Information-Theoretic Analysis of Category Maps and Target Preservation

**DOI:** 10.64898/2026.05.01.722196

**Authors:** Christoph D. Dahl

## Abstract

Categorisation is often treated as a form of compression: a high-dimensional stimulus space is reduced to a smaller set of behaviourally or cognitively useful classes. However, compression alone does not determine whether a category map is useful. The present manuscript develops an information-theoretic framework for evaluating categorisation in terms of both category complexity and target-relevant information preservation. Across a set of synthetic demonstrations, alternative category maps over the same stimulus space are shown to preserve different target variables, including identity, action, nuisance, and hierarchical category structure. The framework is then extended to learned visual representations by analysing layer-derived category maps from a pretrained ResNet-50 network applied to CIFAR-10 images. Two scenarios are compared: a clean-only object run and a pooled nuisance run containing clean, blurred, pixelated, and noise-perturbed images. The results show that category maps can have substantial entropy while preserving information about a variable that is not aligned with the specified target, and that the value of a categorisation depends on the target variable to be preserved. The manuscript argues that categorisation should therefore be evaluated not only by compression or separability, but by the information retained about a specified cognitive, behavioural, or computational target.

## 1 Introduction

Categorisation is one of the central operations of cognition [1, 2]. Organisms and artificial systems encounter a world whose perceptual states are highly variable, continuous, and often noisy. Nevertheless, behaviour rarely depends on every distinguishable detail of that world. Different visual views can be treated as the same object, different individuals can be treated as members of the same kind, and different objects can be grouped according to similar consequences for action. In this broad sense, categorisation compresses the stimulus space: many possible sensory states are mapped onto a smaller number of classes. This compression view is intuitively attractive, but it is incomplete. A compression can be simple without being useful. A stimulus set can be compressed into one category, into arbitrary clusters, or into categories based on irrelevant features. Such category maps reduce complexity, yet they may preserve little information about what the system must recognise, predict, or control. Conversely, a more complex category map may be useful if it preserves a variable that is relevant for behaviour or inference [3, 4, 5]. Thus, the relevant question is not whether categorisation reduces stimulus complexity, but which target-relevant information remains available after that reduction. The importance of this distinction is visible in classical work on categories and concepts. Natural categories are not merely arbitrary partitions of sensory space. They are structured by family resemblance, typicality, and category-level regularities that support inductive generalisation [6, 7, 8, 2]. The basic-level category literature is particularly relevant because it treats category level as a trade-off between informativeness and economy. A basic category such as “dog” or “chair” is often more useful than a very specific subordinate category or a very broad superordinate category because it preserves many predictive regularities while avoiding unnecessary detail [7, 3, 4]. From the present perspective, this can be reformulated as a complexity–relevance problem: different category levels impose different compressions on the same stimulus space, and their usefulness depends on which target variables are preserved. A similar issue appears in exemplar, rational, and computational accounts of concept learning. Categories support generalisation because they encode assumptions about which observations belong together and which properties are expected to transfer across instances [9, 10, 5, 11, 12, 13, 14]. Under this view, a category is not only a label but also a hypothesis about structure in the world. However, different structures can be imposed on the same stimulus set. One partition may preserve identity, another may preserve action relevance, another may preserve context, and another may preserve only nuisance variation. The same sensory observations can therefore support multiple candidate category maps, only some of which are useful for a given inferential problem. The same issue is also relevant to animal categorisation. Non-human animals can learn to sort complex stimuli into experimenter-defined categories and can generalise these discriminations to novel exemplars [15, 16, 17, 18]. These findings are important for the present argument because they show that categorisation can be studied as a relation among stimulus structure, learned partitions, and generalisation targets, without assuming that the category labels correspond to verbal concepts.

Information theory provides a natural language for this issue. Since Shannon [19], entropy has provided a formal measure of uncertainty, and mutual information has quantified how much uncertainty about one variable is reduced by observing another [20]. In sensory coding, efficient coding theories have long treated perceptual systems as transformations that reduce redundancy while preserving behaviourally important information [21]. The information bottleneck framework makes this trade-off explicit by seeking compressed representations that retain information about a relevant target variable [22, 23]. The present approach also builds on the more general idea that transformations can be analysed as information-funnelling operations, in which many initial states are compressed into fewer structured outcomes while entropy and residual state information are tracked across the transformation [24]. Related information-theoretic approaches have been used to analyse behavioural sequences, animal movement patterns, and spatial relations in collective behaviour [25, 26, 27]. In this context, the geometry of animal groups has also been analysed as an information-theoretic problem [28]. The central logic is directly applicable to categorisation. A category map should not be evaluated only by the entropy of the categories it produces. It should be evaluated by the information that those categories preserve about a target variable. This distinction is also relevant for contemporary representation learning. Deep neural networks transform images into hierarchical activation spaces in which object categories, textures, nuisance variables, and task-relevant distinctions may be represented to different degrees [29, 30, 31, 32]. These models have become useful tools for comparing artificial and biological visual representations [33, 34, 35, 36]. Yet the same caution applies: a low-dimensional or clustered representation is not necessarily a semantically useful one [37, 38]. If the dominant structure in an activation space corresponds to blur, noise, or other nuisance variables, an unsupervised category map may preserve those variables more strongly than object identity. In such a case, the representation is compressed and structured, but not useful for the intended semantic target.

The present manuscript develops this point in a deliberately simple formal setting. A category map is defined as a function that assigns each stimulus to a category. Its complexity is measured by the entropy of the induced category variable. Its relevance is measured by the mutual information between the category variable and a target variable. This formulation allows the same stimulus space to be analysed under different candidate target variables, such as identity, action, nuisance condition, or category level. It also allows different category maps to be compared even when they have similar complexity. Four synthetic demonstrations are used to establish the principle. First, several category maps are compared over the same two-dimensional stimulus space, showing that category entropy alone cannot distinguish useful from useless partitions. Second, the same category maps are evaluated with respect to multiple target variables, showing that a category map can be useful for one target and useless for another. Third, robustness under perturbation is examined, showing how preserved information changes when sensory variation increases. Fourth, hierarchical category maps are analysed, showing how subordinate, basic, and broader action/superordinate-like category maps can be interpreted as different compressions of the same stimulus space. A fifth, exploratory demonstration extends the analysis to a pretrained visual neural network. CIFAR-10 images are passed through ResNet-50, activation spaces are clustered at selected layers, and the resulting layer-derived category maps are evaluated against object-relevant and nuisance-relevant targets. This extension is not intended as a full theory of deep-network categorisation. Its purpose is narrower: to show that the same information-theoretic problem arises in learned representations. A non-trivial category map can preserve substantial information, but that information may concern a nuisance variable rather than the semantic target of interest.

## 2 Methods

### 2.1 Overview of the modelling strategy

The analyses were designed to evaluate category maps as target-preserving compressions. A category map was treated as a transformation from a stimulus space to a category variable. The central question was not whether a transformation reduced the complexity of the stimulus space, but whether the resulting categories preserved information about a specified target variable. This logic follows the general information-theoretic distinction between compression and preserved target information [19, 20, 22, 23]. This logic was applied in two stages. First, synthetic stimulus spaces with known generative structure were used to establish the formal relations among category entropy, target information, robustness, and classification level. Second, the same information-theoretic analysis was applied to layer-derived category maps from a pretrained visual neural network. The synthetic demonstrations were deliberately simple. Their purpose was to isolate the formal logic of target-preserving compression, not to model the full complexity of natural category learning. The neural-network extension served a different role. It examined whether the same analysis could be applied to learned representations in a pretrained visual classifier [39, 32], using CIFAR-10 images as a compact natural-image stimulus set [40]. The analysis asked whether unsupervised partitions of activation space preserved object-relevant information or information about nuisance condition, defined here as the image-transformation state: clean, blurred, pixelated, or noise-perturbed. An overview of the five demonstrations, including their stimulus sources, target variables, candidate category maps, and main information-theoretic quantities, is given in Table 1.

**Table 1:**
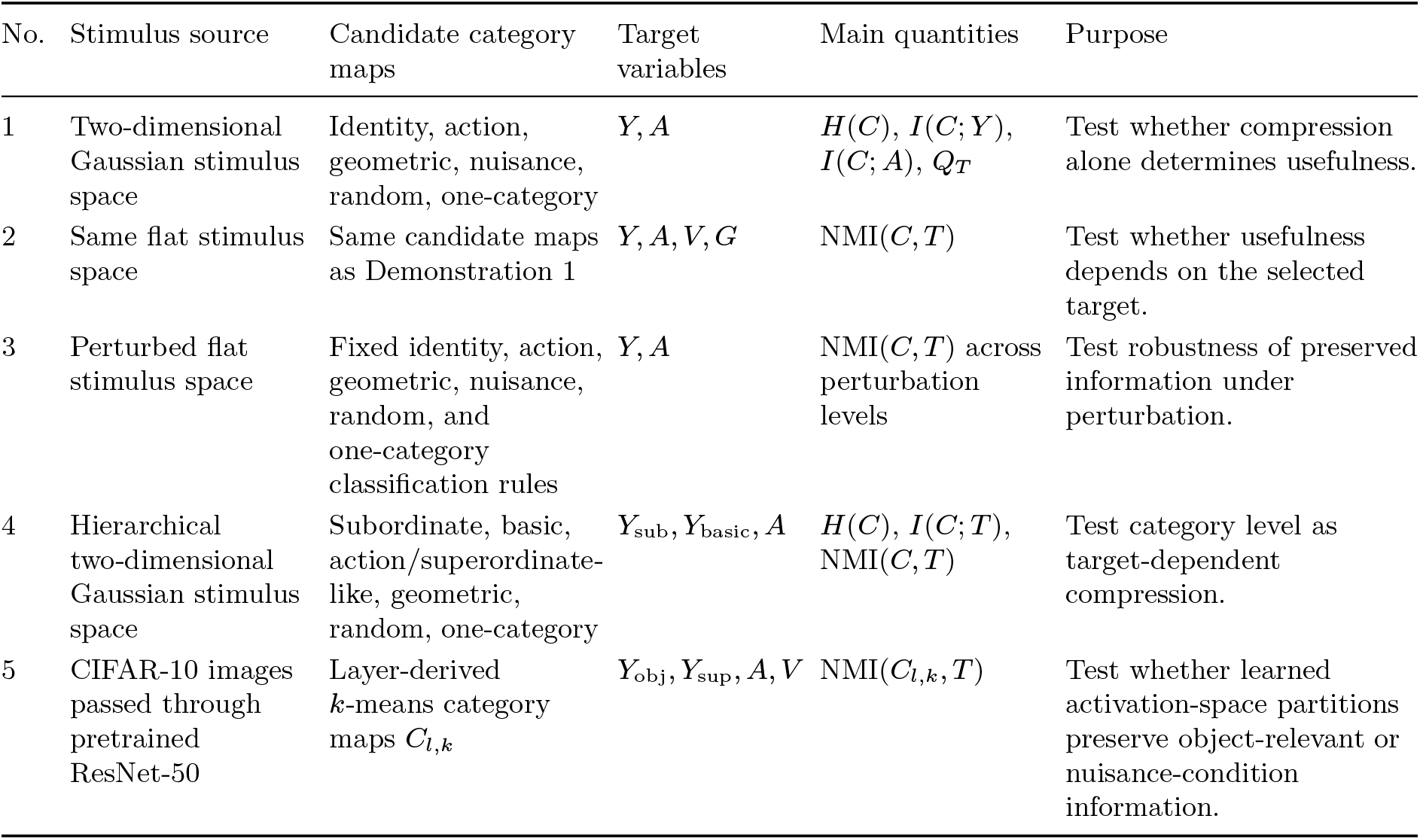
Overview of the five demonstrations. The table summarises the stimulus source, candidate category maps, target variables, main information-theoretic quantities, and purpose of each demonstration.

### 2.2 Category maps as target-preserving compressions

Let *S* denote the stimulus variable, with individual stimulus instances *s* ∈ *S*, and let *C* = *f* (*S*) denote the category variable induced by a category map

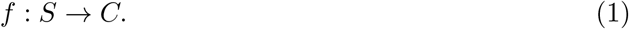

Here, *C* is the category label assigned to each stimulus instance. The category map compresses the stimulus space whenever multiple distinct stimuli are assigned to the same category. All probabilities in the following expressions were estimated from empirical frequencies over the simulated or analysed samples. The complexity of the resulting category variable was quantified by category entropy,

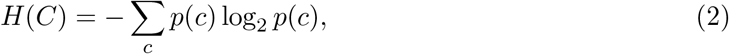

where *p*(*c*) denotes the probability of category *c*. Low entropy indicates a coarse or collapsed category map; high entropy indicates a richer set of category distinctions [19, 20]. Category entropy alone does not specify what information is preserved by the category map. Therefore, each category map was evaluated relative to one or more target variables *T*. The target variable *T* denotes the variable whose preservation is being evaluated, such as identity, action relevance, nuisance condition, or hierarchical category level. Preserved target information was quantified by mutual information,

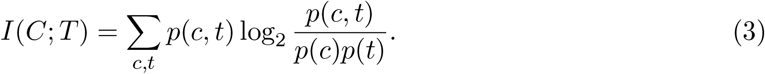

Here, *p*(*c, t*) denotes the joint probability of category *c* and target value *t*, and *p*(*c*) and *p*(*t*) denote the corresponding marginal probabilities. Equivalently,

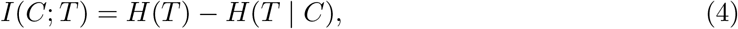

where *H*(*T* | *C*) is the remaining uncertainty about the target after the category is known. For targets with different baseline entropies, preserved information was expressed as normalised mutual information,

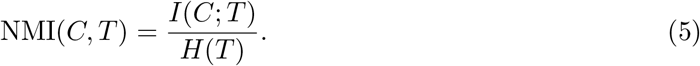

This value measures the fraction of target uncertainty removed by the category map. A value of zero indicates that the category map preserves no information about the target. A value of one indicates that the target is fully determined by the category map. An illustrative quality score was also used to express the trade-off between target preservation and category complexity:

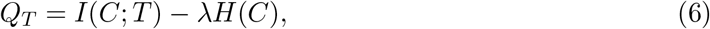

where *λ* controls the strength of the penalty on category complexity. This score follows the same general logic as compression–relevance trade-offs in information-theoretic representation learning, where compressed variables are evaluated by the information they retain about a relevant target [22, 23]. It was not treated as a unique normative measure. Its role was to make the compression–relevance trade-off explicit.

### 2.3 Synthetic demonstrations

#### 2.3.1 Stimulus space and target variables

The synthetic demonstrations used two-dimensional stimulus spaces constructed from Gaussian clusters. Each point represented one stimulus instance *s*_*i*_ with two continuous feature values,

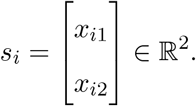

The feature space was deliberately minimal, because the aim was not to model natural category learning in full detail, but to isolate the formal logic of target-preserving compression under controlled conditions. In this setting, the stimulus variable *S* denotes the set of sampled stimulus points,

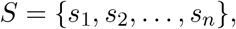

and each candidate category map assigns every stimulus point *s*_*i*_ to a category label *C*_*i*_. The same stimulus set was assigned multiple target variables. In the flat stimulus-space demonstrations, four targets were defined:

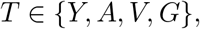

where *Y* denotes an identity-like target, *A* an action-like target, *V* a nuisance target, and *G* a geometric target. The identity target *Y* indicated the latent cluster or individual-like class from which a stimulus was generated. The action target *A* grouped stimuli according to a shared consequence or response-relevant class. The nuisance target *V* represented a variable that could structure the stimulus set without being relevant to identity or action. The geometric target *G* described a partition based on spatial organisation in the two-dimensional feature space. These targets were not intended as empirical claims about actual visual categories. They served as controlled variables that made it possible to ask how the same category map can preserve different kinds of information. This use of multiple possible targets follows the general idea that categories support different forms of generalisation depending on which structure is treated as relevant [9, 10, 5, 12]. The demonstrations therefore separate the existence of a category partition from the question of which target variable that partition preserves. For each synthetic demonstration, stimulus points were sampled from Gaussian clusters with fixed means and covariance structures, with equal numbers of samples drawn per latent cluster. Unless otherwise noted, the same sampled stimulus set was then evaluated under alternative target assignments and category maps. In the synthetic demonstrations, the same low-dimensional stimulus space was evaluated under several alternative target assignments and category maps. In the flat demonstrations, the sampled stimuli were analysed relative to identity-like, action-like, nuisance, and geometric targets. Candidate category maps were then defined to preserve identity, action, geometric, nuisance, random, or fully collapsed one-category structure. In the perturbation analysis, Gaussian noise was added to the stimulus coordinates before re-evaluating fixed baseline category-assignment rules. In the hierarchical demonstration, a separate stimulus space was constructed such that subordinate, basic-level, and broader action/superordinate-like targets could all be defined over the same points. This arrangement allowed category level to be treated explicitly as a target-dependent compression of the same underlying stimulus space.

#### 2.3.2 Candidate category maps

Candidate category maps were defined over the same stimulus space. Each map *m* induced a category variable *C*^(*m*)^ by assigning every stimulus instance to a category label:

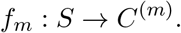

The set of candidate maps was

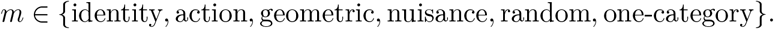

The identity-preserving map preserved the latent identity distinctions *Y*. The action-preserving map merged identities that shared the same action consequence *A*. The geometric clustering map grouped stimuli according to their spatial arrangement in feature space. The nuisance-based map grouped stimuli according to a context or nuisance variable *V*. The random partition imposed labels independent of the relevant target variables. The one-category map assigned all stimuli to a single category and therefore represented maximal compression,

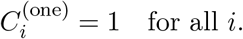

Thus, each category map produced a category variable *C*^(*m*)^, but the meaning of that category variable depended on the map that generated it. Some maps were aligned with a target variable, whereas others were deliberately misaligned. This distinction is important because category complexity alone does not imply target usefulness. A category map can have several categories and therefore non-zero entropy *H*(*C*^(*m*)^), while preserving little or no information about the target variable of interest. These maps were evaluated against the defined target variables using category entropy, mutual information, conditional entropy, and normalised mutual information. Category entropy *H*(*C*^(*m*)^) quantified the complexity of the category labels produced by map *m*. Mutual information *I*(*C*^(*m*)^; *T*) quantified how much information the map preserved about a target variable *T*. Conditional entropy *H*(*T* | *C*^(*m*)^) quantified how much target uncertainty remained after the category label was known. Normalised mutual information was computed as

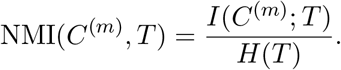

This expressed preserved information as the fraction of target uncertainty removed by the category map. The notation *C*^(*m*)^ is used here to emphasise that the category variable depends on the selected map *m*. This allowed category complexity and target preservation to be separated explicitly [19, 20, 22, 23].

#### 2.3.3 Demonstrations 1–3: flat stimulus space

Demonstration 1 evaluated whether compression alone was sufficient for category usefulness. Several category maps were compared over the same stimulus space. The key comparison was between category complexity, measured by

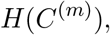

and preserved target information, measured by

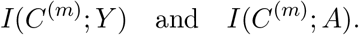

This demonstration tested whether category maps with similar entropy could differ in target relevance. It also connected the formal analysis to earlier work treating category utility as a relation between category structure and informative features [3, 4].

Demonstration 2 evaluated the same category maps against multiple target variables. Each map was compared with identity *Y*, action *A*, nuisance condition *V*, and geometric structure *G*:

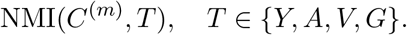

This demonstration tested whether category usefulness was an intrinsic property of a category map or a relation between a map and a specified target variable. In information-theoretic terms, the same category variable *C*^(*m*)^ can have high NMI(*C*^(*m*)^, *T*) for one target *T*, but low NMI(*C*^(*m*)^, *T* ^*′*^) for another target *T* ^*′*^:

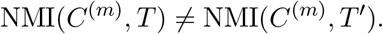

Therefore, usefulness was defined relative to the target that the category system was expected to preserve.

Demonstration 3 examined robustness under sensory perturbation. Noise was added to the stimulus features, producing perturbed stimulus instances

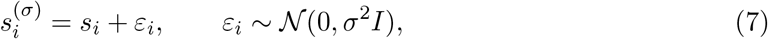

where *σ* denotes perturbation strength. Fixed baseline classification rules were then re-evaluated across perturbation levels. Robustness was quantified as the stability of normalised mutual information under perturbation:

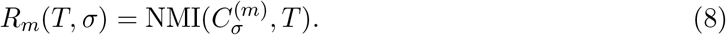

Here, 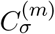 denotes the category variable induced by map *m* when applied at perturbation strength *σ*. A robust target-preserving map should retain relatively high *R*_*m*_(*T, σ*) as the stimulus coordinates become noisier, whereas a non-aligned map should remain uninformative about the target. This demonstration tested whether target-preserving category maps retained relevant information when the stimulus space was degraded.

#### 2.3.4 Demonstration 4: hierarchical category maps

A separate synthetic demonstration introduced a hierarchical stimulus space. Stimuli were organised such that subordinate identity, basic class, and broader action/superordinate-like targets could all be defined over the same points:

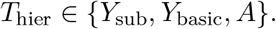

Candidate category maps were constructed at corresponding levels:

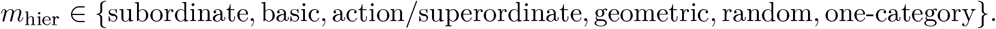

This demonstration tested whether classification level could be interpreted as a target-dependent compression. A subordinate category map preserves fine identity distinctions but has higher category complexity. A basic-level category map discards some subordinate distinctions while preserving broader class structure. An action/superordinate-like map compresses further and preserves still broader target structure. This formulation is closely related to the classical idea that category levels differ in their balance between informativeness and economy [7, 6, 2]. In the present analysis, however, this balance was expressed directly in terms of

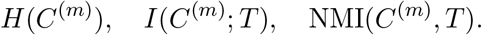

Thus, subordinate, basic, and action/superordinate-like classifications were treated as alternative compressions of the same stimulus space, whose usefulness depended on which target variable was to be preserved.

### 2.4 Neural-network extension

#### 2.4.1 Image dataset and target variables

To examine whether the same information-theoretic analysis could be applied to learned visual representations, an exploratory neural-network demonstration was carried out using a pretrained ResNet-50 model [39]. ResNet-50 was used here as a fixed feature extractor, not as a model retrained for the present task. CIFAR-10 images [40] were used as a compact natural-image stimulus set. This extension follows the broader use of deep neural networks as models of hierarchical visual representation and as comparison systems for biological object recognition [29, 33, 36, 32].

Each image *s*_*i*_ was assigned to four target variables:

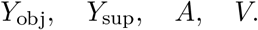

The object-class target *Y*_obj_ corresponded to the ten CIFAR-10 categories:

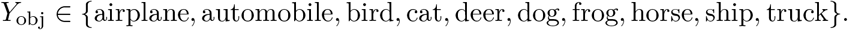

These CIFAR-10 labels are treated here as basic-level-like object classes rather than as subordinate categories. The superordinate target *Y*_sup_ distinguished animals from vehicles:

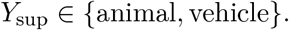

The environment- or affordance-like target *A* distinguished air-, land-, and water-associated categories:

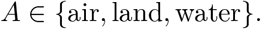

The nuisance target *V* denoted the image-transformation condition:

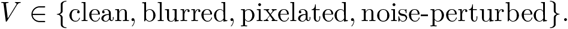

Here, *V* was treated as a nuisance variable only relative to object-relevant targets. If the target were to detect image degradation, *V* would be target-relevant. In the present analysis, however, *V* was used to test whether unsupervised activation-space partitions preserved transformation condition rather than object-relevant category structure. Here, *y*_*i*_ denotes the original CIFAR-10 label of image *s*_*i*_. The mapping from CIFAR-10 labels to the superordinate target was defined as follows:

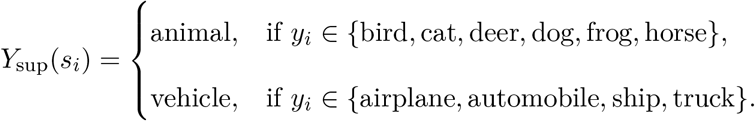

The environment- or affordance-like target was defined as

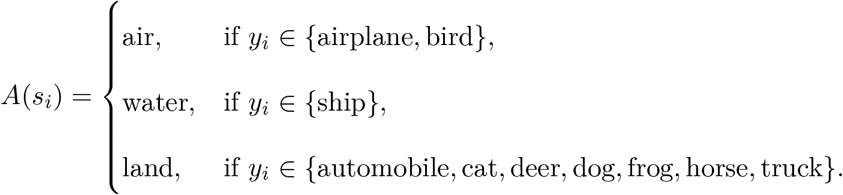

This target was not treated as a literal action label. It was included as a constructed coarse consequence-relevant grouping, allowing the same layer-derived category maps to be evaluated against a target that was broader than the CIFAR-10 object-class target but different from the animal–vehicle distinction.

#### 2.4.2 Clean-only and pooled nuisance scenarios

Two ResNet-50 scenarios were analysed. In the clean-only object run, 300 images from each CIFAR-10 class were selected, producing 10 × 300 = 3000 clean images. This scenario tested whether unsupervised activation-space partitions preserved object-relevant information when systematic nuisance variation was absent. In the clean-only run, *V* was degenerate because all images belonged to the clean condition. Nuisance information was therefore treated as zero by convention and was used only as a reference baseline, not as a meaningful normalised target. In the pooled nuisance run, the same number of original images was selected, but each image was represented in four versions: clean, blurred, pixelated, and noise-perturbed. This produced 10 × 300 × 4 = 12000 exported images, from which 3000 images were selected for the activation analysis. The activation analysis was limited to 3000 images for computational comparability with the clean-only run. This subset was sampled after image generation and was stratified by object class *Y*_obj_ where possible. This scenario tested whether the same unsupervised procedure preserved object-relevant information or instead captured transformation-induced nuisance structure when image condition varied systematically. The contrast between the two scenarios was therefore designed to distinguish object-relevant preservation from nuisance-condition preservation.

### 2.4.3 Network model and layer activations

For each selected ResNet-50 layer *l*, an activation vector was extracted for each image:

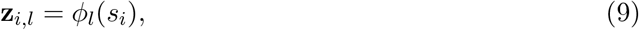

where *s*_*i*_ denotes image *i, ϕ*_*l*_(*·*) denotes the transformation implemented by ResNet-50 up to layer *l*, and **z**_*i,l*_ denotes the corresponding activation vector. Six layers were selected to sample early, intermediate, late, and final network representations:

~~~
activation_1_relu, activation_10_relu, activation_22_relu,
activation_40_relu, activation_49_relu, avg_pool.
~~~

These layers were selected to sample the representational hierarchy from early convolutional processing to the final pooled representation before classification. The layer-derived activation spaces were treated as learned representational spaces in which object category, broader category structure, affordance-like structure, and nuisance condition could be preserved to different degrees [35, 32, 38]. Before clustering, activation vectors were standardised and reduced by principal component analysis. Thirty principal components were retained, yielding a reduced feature vector

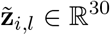

for each image and layer. For each scenario, layer, and value of *k*, clustering was applied to the standardised PCA-reduced activation matrix. *k*-means was run with five replicates and a maximum of 200 iterations per replicate. The random seed was fixed at 1 to make image selection and clustering reproducible. The reduced activation space was then partitioned by *k*-means clustering. For each layer *l* and number of clusters *k*, this produced a layer-specific category map

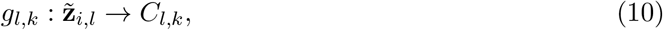

where *C*_*l,k*_ denotes the cluster label assigned to image *i*. Thus, *C*_*l,k*_ was treated as the category variable induced by unsupervised clustering of the activation geometry at layer *l*. Analyses were carried out for

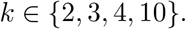

These values were selected because they correspond to the number of levels in the main target variables: two superordinate classes in *Y*_sup_, three environment- or affordance-like classes in *A*, four nuisance conditions in *V*, and ten object classes in *Y*_obj_. The main manuscript figure focuses on *k* = 10, because this is the richest tested category map and allows the strongest comparison between object-class, superordinate, affordance-like, and nuisance-condition preservation. The additional *k*-values were retained in the analysis pipeline as diagnostic checks of how preserved target information changed with category-map granularity.

#### 2.4.4 Information-theoretic evaluation of layer-derived category maps

For each layer *l* and each value of *k*, the entropy of the induced category variable *C*_*l,k*_ was computed:

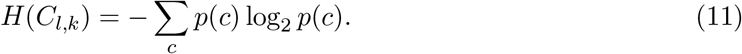

This value quantified the complexity of the unsupervised layer-derived category map. The same category map was then evaluated against each target variable:

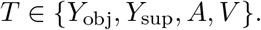

Preserved target information was quantified by

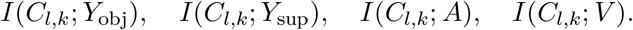

The corresponding normalised values were computed as

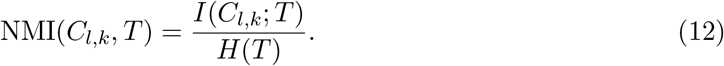

Thus, NMI(*C*_*l,k*_, *T*) measured the fraction of uncertainty about target *T* that was removed by knowing the layer-derived cluster label *C*_*l,k*_. A value close to zero indicated that the cluster labels preserved little information about the target. A value close to one indicated that the cluster labels almost fully determined the target. This analysis quantified whether a layer-derived category map preserved object-class information, superordinate animal–vehicle information, environment- or affordance-like information, or nuisance-condition information. Importantly, the procedure did not train a supervised classifier for each target. Instead, it asked which target variables were already preserved by unsupervised partitions of the activation space. This distinction was central to the neural-network extension: a layer-derived category map could be structured and non-trivial, but still preserve information primarily about nuisance condition rather than the intended object-relevant target.

### 2.5 Implementation and reproducibility

All simulations and neural-network analyses were implemented in MATLAB. Synthetic demonstrations generated the stimulus spaces, target variables, candidate category maps, information-theoretic quantities, and figures directly from the analysis scripts. The neural-network extension used CIFAR-10 image batches [40], a pretrained ResNet-50 model [39], principal component analysis, *k*-means clustering [41], and the same entropy and mutual-information functions used for the synthetic demonstrations [19, 20]. Because the synthetic demonstrations used known generative structures and explicitly defined category maps, the main comparisons were information-theoretic rather than inferential. The reported values quantify the preservation properties of each category map directly, rather than estimating population effects from sampled participants or empirical trials. The neural-network extension was treated as an exploratory computational demonstration of the same principle in learned representations.

## 3 Results

### 3.1 Demonstration 1: compression alone is insufficient

The first demonstration evaluated six candidate category maps over the same two-dimensional stimulus space (Figure 1). The stimulus coordinates were held constant across panels. Only the assignment of points to categories was changed. The identity-preserving, action-preserving, and geometric maps imposed partitions that were aligned with different target structures, whereas the nuisance-based and random maps imposed non-trivial partitions that were not aligned with the identity or action targets. The one-category map represented maximal compression by assigning all stimuli to a single class. The visual comparison in Figure 1 shows why the geometry of the stimulus space is not sufficient to define a category map. The same three-cluster distribution can be partitioned as an identity map, an action map, a geometric map, a nuisance map, a random map, or a collapsed one-category map. These alternatives differ not because the stimuli differ, but because different variables are preserved by the mapping from stimuli to categories. The information-theoretic summaries in Figure 2A–D show that category entropy alone was insufficient to determine category usefulness, consistent with the distinction between category complexity and preserved target information [19, 20, 3, 4]. In Figure 2A, the identity-preserving map had high category entropy and preserved the identity target almost fully, whereas the action-preserving map had lower category entropy and preserved the action target. The nuisance-based and random maps retained category complexity, but preserved almost no information about the identity or action targets. Conversely, the one-category map achieved maximal compression while leaving target uncertainty largely unchanged. Thus, a category map could be complex without being useful, and a category map could be simple without preserving the target of interest. The quality-score and conditional-entropy panels make the same point from complementary directions. In Figure 2B, penalising category complexity favoured category maps that preserved the selected target rather than merely producing many categories. Figure 2C shows that preserved target information did not follow category entropy alone: maps with similar entropy differed in the information they retained about identity or action. In Figure 2D, conditional entropy showed the residual uncertainty after categorisation: target-aligned maps reduced uncertainty about the corresponding target, whereas nuisance-based, random, and one-category maps left substantial identity or action uncertainty. Therefore, compression was not sufficient. Relevant information had to be preserved by the compression.

**Figure 1:**
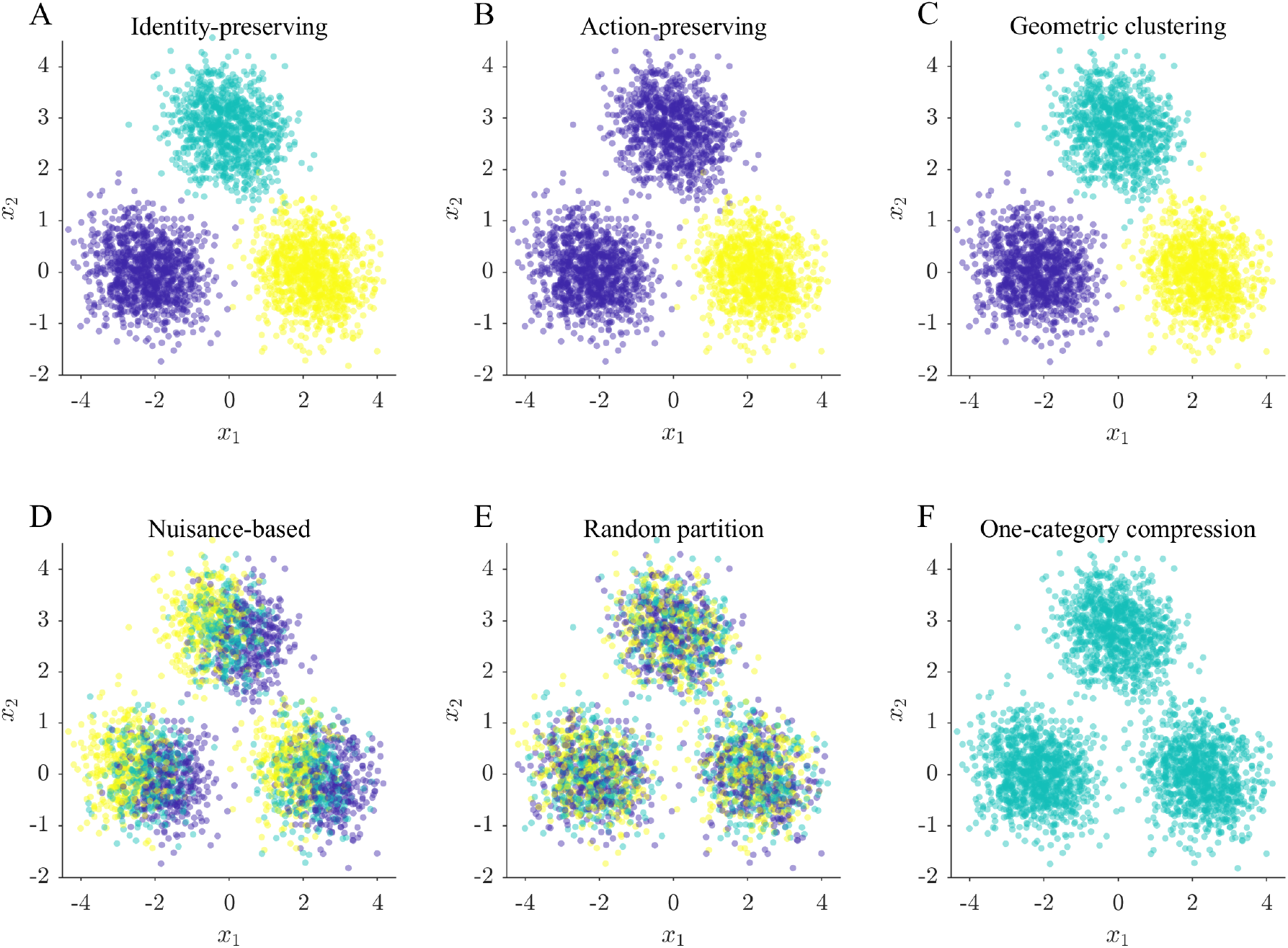
Candidate category maps over the same two-dimensional stimulus space. Panels show (A) identity-preserving, (B) action-preserving, (C) geometric clustering, (D) nuisance-based, (E) random partition, and (F) one-category compression maps. The stimulus coordinates are identical across panels; only the category assignment changes. The figure illustrates that the same stimulus geometry can support several possible compressions, only some of which preserve the target variable of interest.

**Figure 2:**
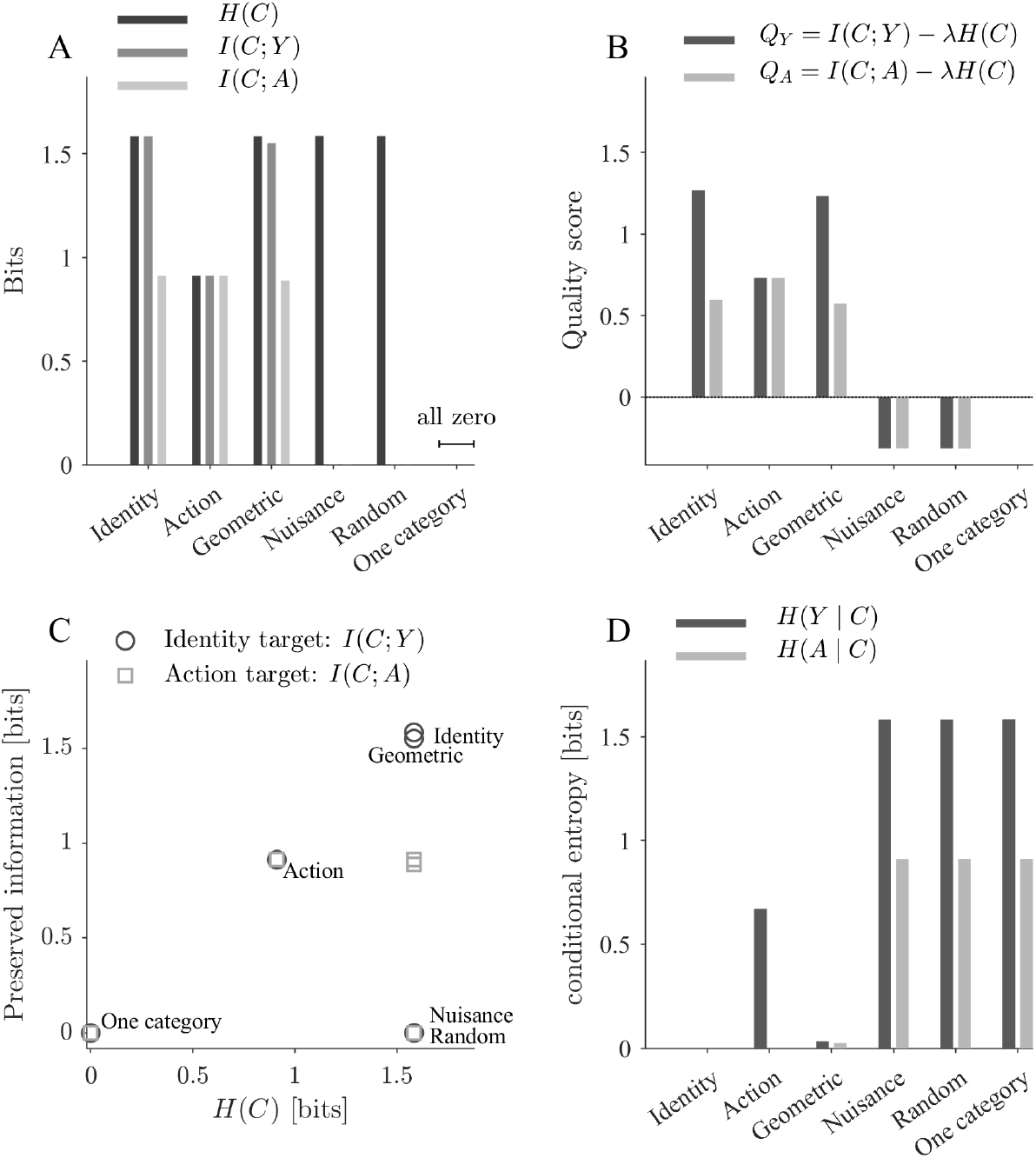
Information-theoretic comparison of the candidate category maps. (A) Category entropy *H*(*C*), identity information *I*(*C*; *Y*), and action information *I*(*C*; *A*). (B) Target-specific quality scores *Q*_*Y*_ = *I*(*C*; *Y*) *λH*(*C*) and *Q*_*A*_ = *I*(*C*; *A*) *λH*(*C*). (C) Preserved target information plotted against category entropy. Circles indicate the identity target *I*(*C*; *Y*) and squares indicate the action target *I*(*C*; *A*); text labels indicate the candidate category maps. (D) Conditional entropy after categorisation, showing the remaining identity uncertainty *H*(*Y C*) and action uncertainty *H*(*A C*). Together, the panels show that category complexity alone is insufficient: useful compression requires preservation of the selected target variable.

### 3.2 Demonstration 2: preserved information depends on the target

The second demonstration evaluated the same candidate category maps against four target variables: identity *Y*, action *A*, nuisance condition *V*, and geometric structure *G* (Figure 3A,B). Figure 3A shows the resulting target-by-map matrix of normalised mutual information, whereas Figure 3B presents the same values as grouped bars. The result showed that no category map was intrinsically useful in a target-independent sense. Rather, usefulness depended on the relation between the category map and the variable that was to be preserved, consistent with target-dependent accounts of compression and category utility [3, 4, 22, 23]. The identity-preserving map preserved the identity target fully and, because of the construction of the stimulus space, also preserved the action target. It additionally preserved most of the geometric target. The action-preserving map fully preserved the action target, but preserved only part of the identity and geometric targets. The geometric map preserved the geometric target and also preserved much of the identity and action structure because the geometric clusters were aligned with the underlying stimulus clusters. The nuisance map preserved the nuisance target, but preserved essentially none of the identity, action, or geometric targets. The random and one-category maps preserved no target information in this construction. Together, Figure 3A and B make the target-dependence of category usefulness explicit. A nuisance category is useful if nuisance condition is the target, but useless if object identity, action relevance, or geometric structure is the target. Likewise, a coarse action category can be useful for action selection while being insufficient for subordinate identity recognition. Category maps therefore require an explicit target variable for evaluation; the relevant question is not whether a map compresses the stimulus space, but which variable remains informative after compression.

**Figure 3:**
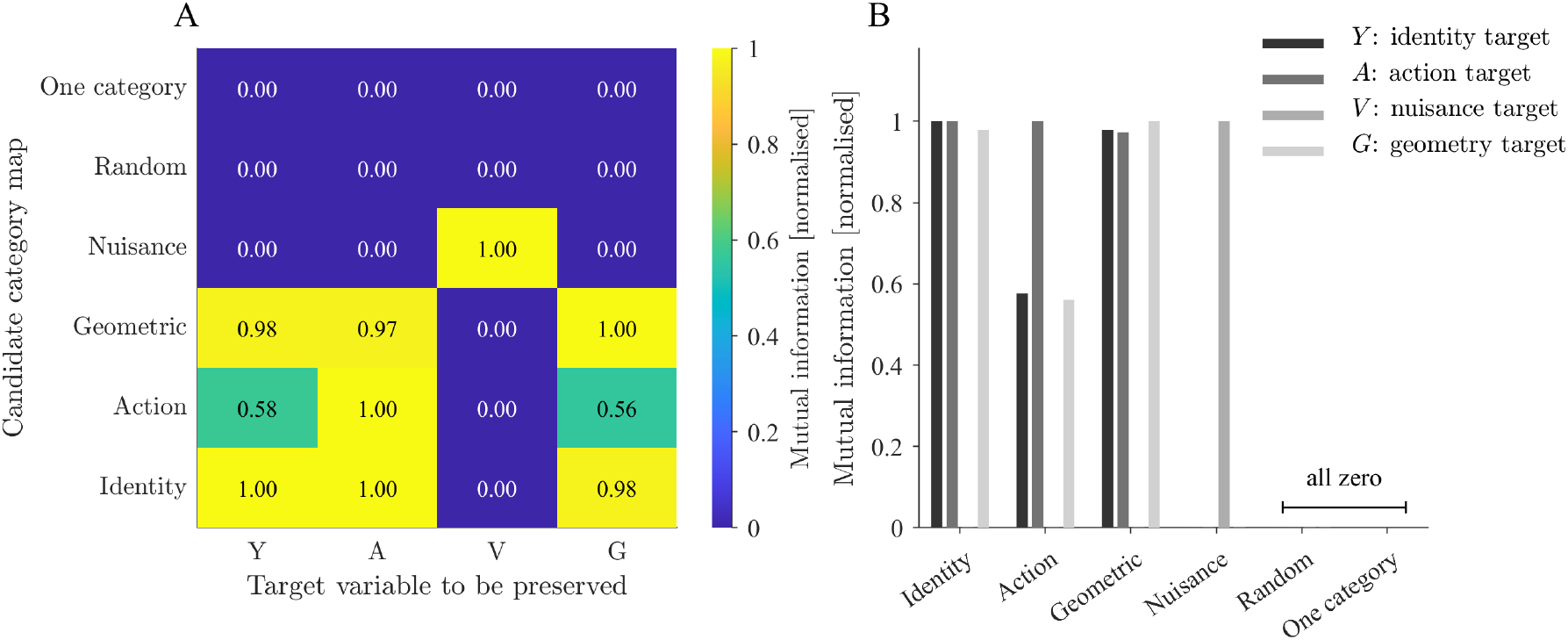
Target-specific preservation by the same candidate category maps. (A) Normalised mutual information between each candidate category map and each target variable. Rows correspond to category maps and columns correspond to target variables: identity *Y*, action *A*, nuisance condition *V*, and geometric structure *G*. (B) The same values shown as grouped bars. The identity, action, geometric, and nuisance maps are informative for different targets, whereas the random and one-category maps preserve no target information in this construction.

### 3.3 Demonstration 3: robustness of preserved information under perturbation

The third demonstration examined how preserved information changed under sensory perturbation, separately for identity-relevant information (Figure 4A) and action-relevant information (Figure 4B). Perturbations were introduced into the stimulus features, and fixed baseline classification rules were applied to the perturbed stimulus coordinates. The analysis asked whether target-aligned classification rules continued to preserve identity-relevant and action-relevant information as sensory noise increased. This question is related to the broader problem of whether representations preserve task-relevant structure while becoming invariant, or at least robust, to nuisance variation [31, 38]. For identity-relevant information, the identity and geometric classification rules produced nearly overlapping robustness profiles (Figure 4A). Both preserved high identity information under weak perturbation, followed by a gradual decline as perturbation strength increased. The action classification rule preserved less identity information, but remained above zero across the tested perturbation range because the action partition retained part of the identity structure. Nuisance, random, and one-category classification rules overlapped at zero, indicating that these classification rules preserved essentially no identity-relevant information. For action-relevant information, the identity, action, and geometric classification rules overlapped closely (Figure 4B). This is expected from the construction of the stimulus space: the identity and geometric distinctions retained the action-relevant grouping, and the action classification rule preserved it directly. As perturbation increased, preserved action information decreased gradually, but remained substantially above the nuisance, random, and one-category classification rules. Thus, robustness depended on alignment between the classification rule and the target variable, not merely on the existence of a category partition.

**Figure 4:**
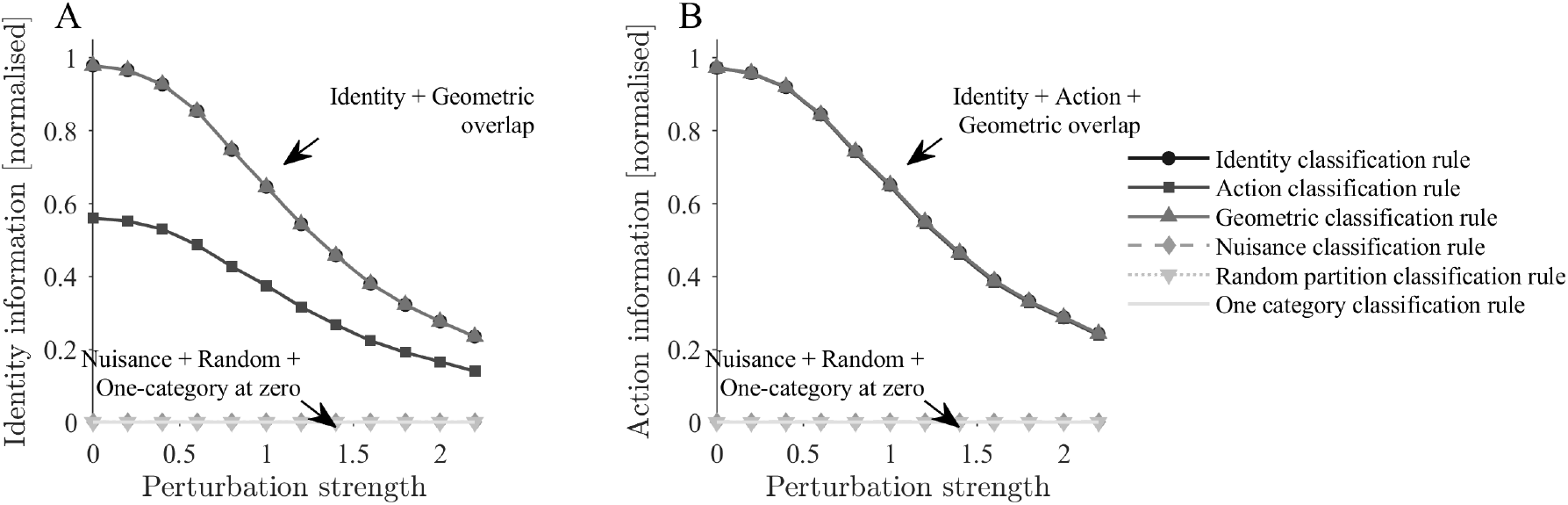
Robustness of preserved target information under sensory perturbation. (A) Normalised identity information as a function of perturbation strength. Identity and geometric classification rules overlap closely, whereas nuisance, random, and one-category classification rules overlap at zero. (B) Normalised action information as a function of perturbation strength. Identity, action, and geometric classification rules overlap closely because all three preserve the action-relevant structure in the present stimulus construction. Nuisance, random, and one-category classification rules again remain at zero.

### 3.4 Demonstration 4: hierarchical category maps over the same stimulus space

The fourth demonstration examined hierarchical category maps over a stimulus space in which subordinate identities, basic-level categories, and a broader action/superordinate-like target were defined over the same points (Figure 5A–F). Candidate maps were constructed at corresponding levels of specificity, together with geometric, random, and one-category alternatives. This allowed category level to be treated as an information-preservation problem rather than as a fixed naming hierarchy. This framing is related to classical work on basic-level categories, in which category levels are understood as differing in their balance between informativeness, generality, and economy [6, 7, 2]. The visual partitions in Figure 5A–F show that the same hierarchical stimulus space can be compressed at different levels. The subordinate map preserved the finest identity distinctions. The basic map merged subordinate identities into broader kinds. The action-level map compressed the space further by preserving the action-relevant grouping. The geometric map recovered much of the basic/action structure because spatial organisation was aligned with these broader targets. The close correspondence between the basic-level map and the geometric clustering map is informative: in this synthetic construction, the basic-level categories coincide with the dominant geometry of the stimulus space. Thus, the geometric map preserves basic-level information not because it was defined by the basic-level labels, but because the basic-level labels are aligned with compact regions of the feature space. The random map retained category complexity without target alignment, whereas the one-category map collapsed all stimuli into a single class. This should not be interpreted as a general claim that basic-level categories are always geometrically obvious; rather, it shows that category level and stimulus-space geometry can coincide when the relevant features are organised in clustered form. In Figure 6A, the subordinate map preserved subordinate information fully, whereas the basic, action, and geometric maps preserved different broader targets. In Figure 6B, the subordinate map preserved the finest target at higher category entropy, the basic map discarded subordinate detail while preserving basic and action-relevant information, and the action map imposed the strongest non-trivial target-aligned compression while preserving the action target. The geometric map preserved basic and action information because of the spatial structure of the synthetic stimulus space. Random and one-category maps were not useful for the defined targets: the random map retained entropy without preserving target information, and the one-category map collapsed both complexity and target preservation. This result shows that category level can be interpreted as a trade-off between complexity and preserved target information. Subordinate, basic, and action/superordinate-like categories are not merely points on a naming hierarchy. They are different compressions of the same stimulus space, each preserving a different target structure. In this sense, category level is target-dependent rather than category-level independent. The same partition may be appropriate for one inferential or behavioural demand and inappropriate for another. This interpretation is consistent with accounts in which category usefulness depends on the information that categories preserve about relevant properties, rather than on category granularity alone [3, 4].

**Figure 5:**
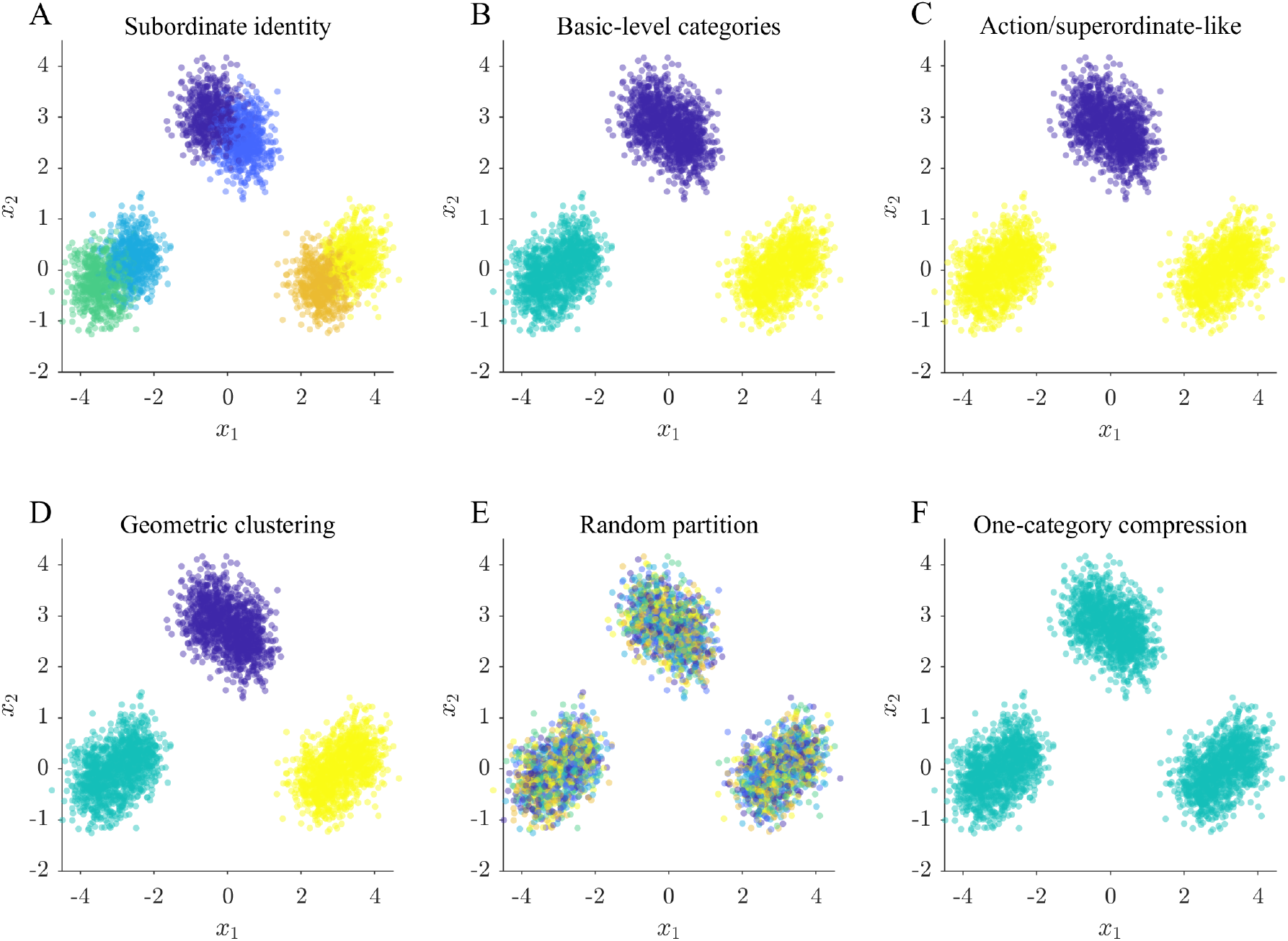
Hierarchical category maps over the same stimulus space. Panels show (A) subordinate identity, (B) basic-level category, (C) action/superordinate-like, (D) geometric clustering, (E) random partition, and (F) one-category compression maps in the hierarchical demonstration. The panels illustrate that subordinate, basic-level, action/superordinate-like, geometric, random, and collapsed maps impose different compressions on the same underlying stimulus distribution.

**Figure 6:**
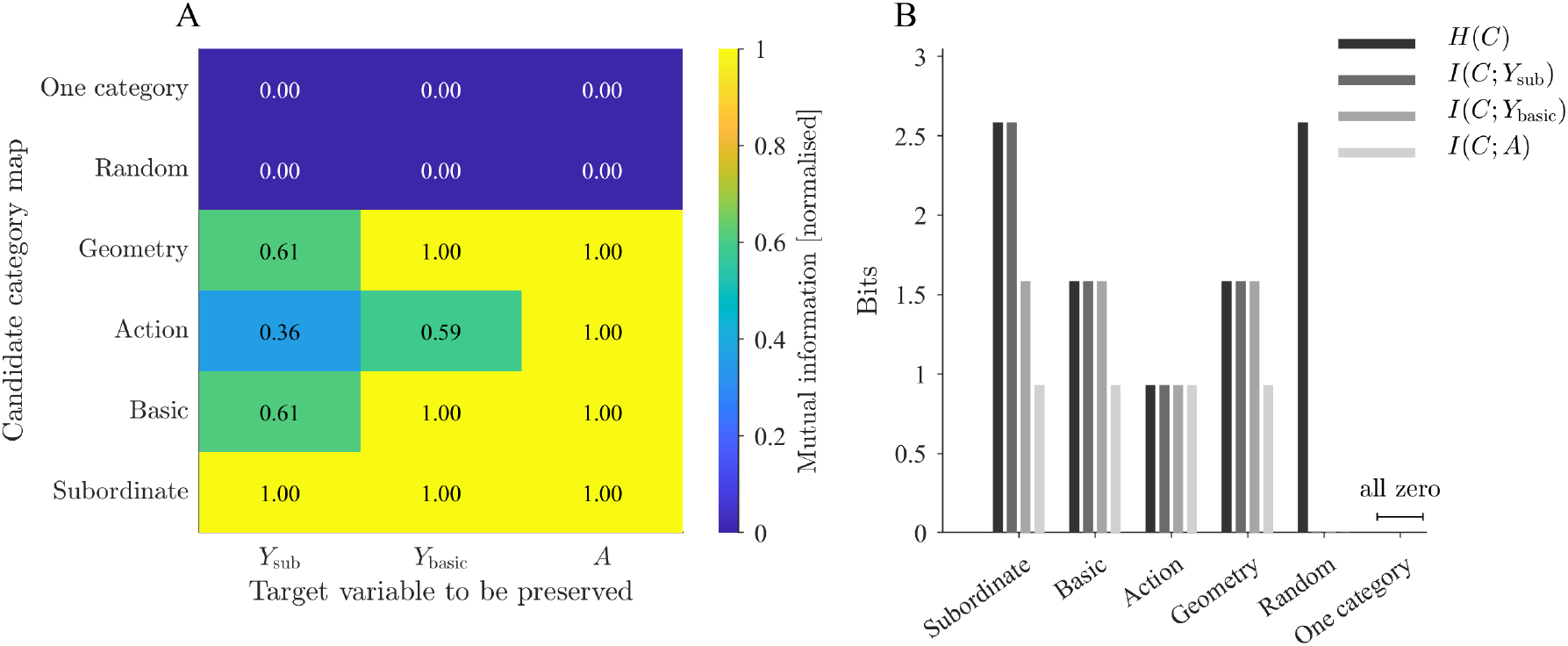
Information-theoretic analysis of hierarchical category maps. (A) Normalised mutual information between each hierarchical category map and each target variable. Rows correspond to candidate maps and columns correspond to target variables. (B) Category entropy and preserved target information for subordinate identity *Y*_sub_, basic-level target *Y*_basic_, and action target *A*. The subordinate map preserves the most fine-grained target at higher complexity, whereas basic and action maps impose stronger compression while preserving broader target variables. The random map retains category entropy without useful target preservation, and the one-category map is an all-zero case.

### 3.5 Demonstration 5: layer-derived category maps in a pretrained visual network

The preceding demonstrations used explicitly constructed category maps over controlled stimulus spaces. The fifth demonstration examined whether the same target-preservation logic could be applied to category maps induced from the activation geometry of a pretrained visual network. ResNet-50 activations were extracted for CIFAR-10 images [39, 40], and unsupervised *k*-means partitions of the reduced activation spaces were treated as layer-derived category maps. This follows the broader use of deep neural networks as hierarchical visual representation models and as comparison systems for biological object recognition [33, 36, 32]. These maps were then evaluated against object class, superordinate animal–vehicle class, environment- or affordance-like class, and nuisance condition. The main network summary is shown for *k* = 10, because this value matches the number of CIFAR-10 object classes and provides the strongest test of whether unsupervised activation-space partitions preserve object-relevant information rather than information about image-transformation condition (Figure 7A–C). Figure 8 provides a qualitative single-example illustration of how the same object, under different nuisance transformations, is represented across the analysed network layers. This visualisation complements the information-theoretic summaries in Figure 7 by showing that transformation-dependent structure remains visible across much of the representational hierarchy.

**Figure 7:**
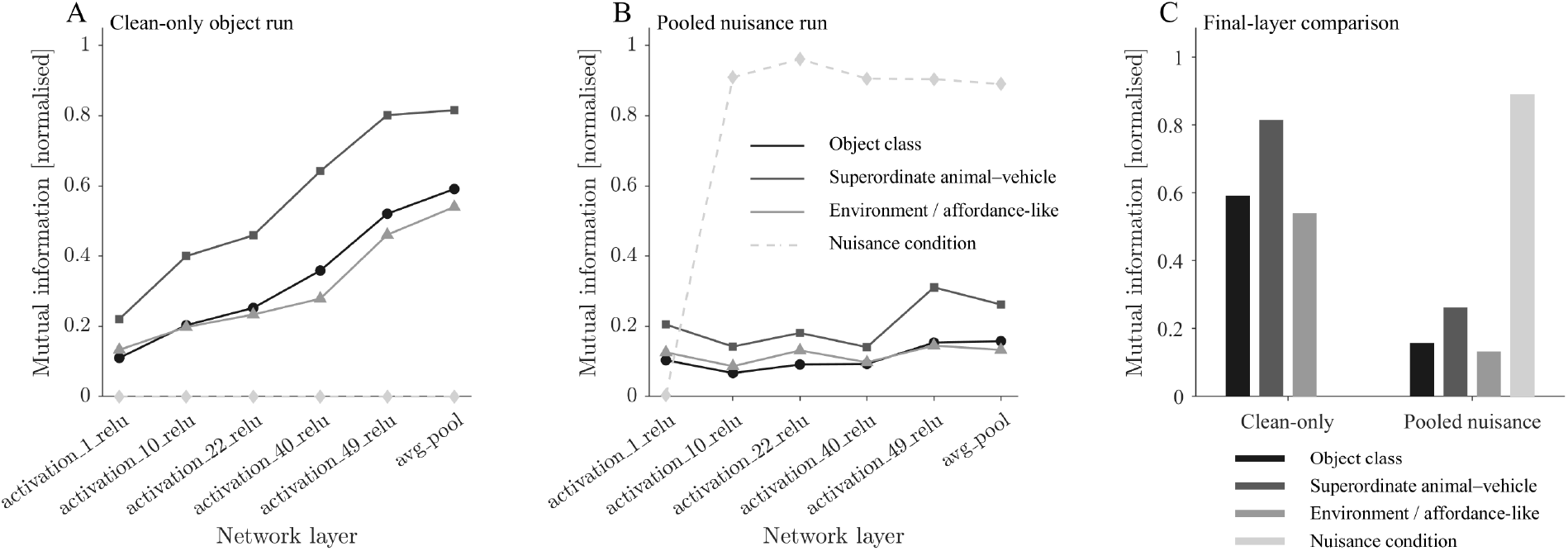
Layer-derived category maps in a pretrained ResNet-50 model. (A) Clean-only object run with *k* = 10: normalised mutual information between the layer-derived category map *C*_*l*,10_ and each target variable across the analysed network layers. Object-relevant targets, especially the superordinate animal–vehicle target, increased across the network hierarchy, whereas nuisance information was absent. (B) Pooled nuisance run with clean, blurred, pixelated, and noise-perturbed images: nuisance condition became the dominant preserved target, while object-class, superordinate animal–vehicle, and environment- or affordance-like information remained comparatively weak. The line legend shown in panel B applies to both panels A and B. (C) Final-layer comparison at `avg_pool`, showing that clean-only partitions preserved object-relevant information, whereas pooled-nuisance partitions preferentially preserved nuisance condition. Panel C uses the bar legend shown below the axis. All values are normalised mutual information values, *I*(*C*_*l*,10_; *T*)*/H*(*T*).

**Figure 8:**
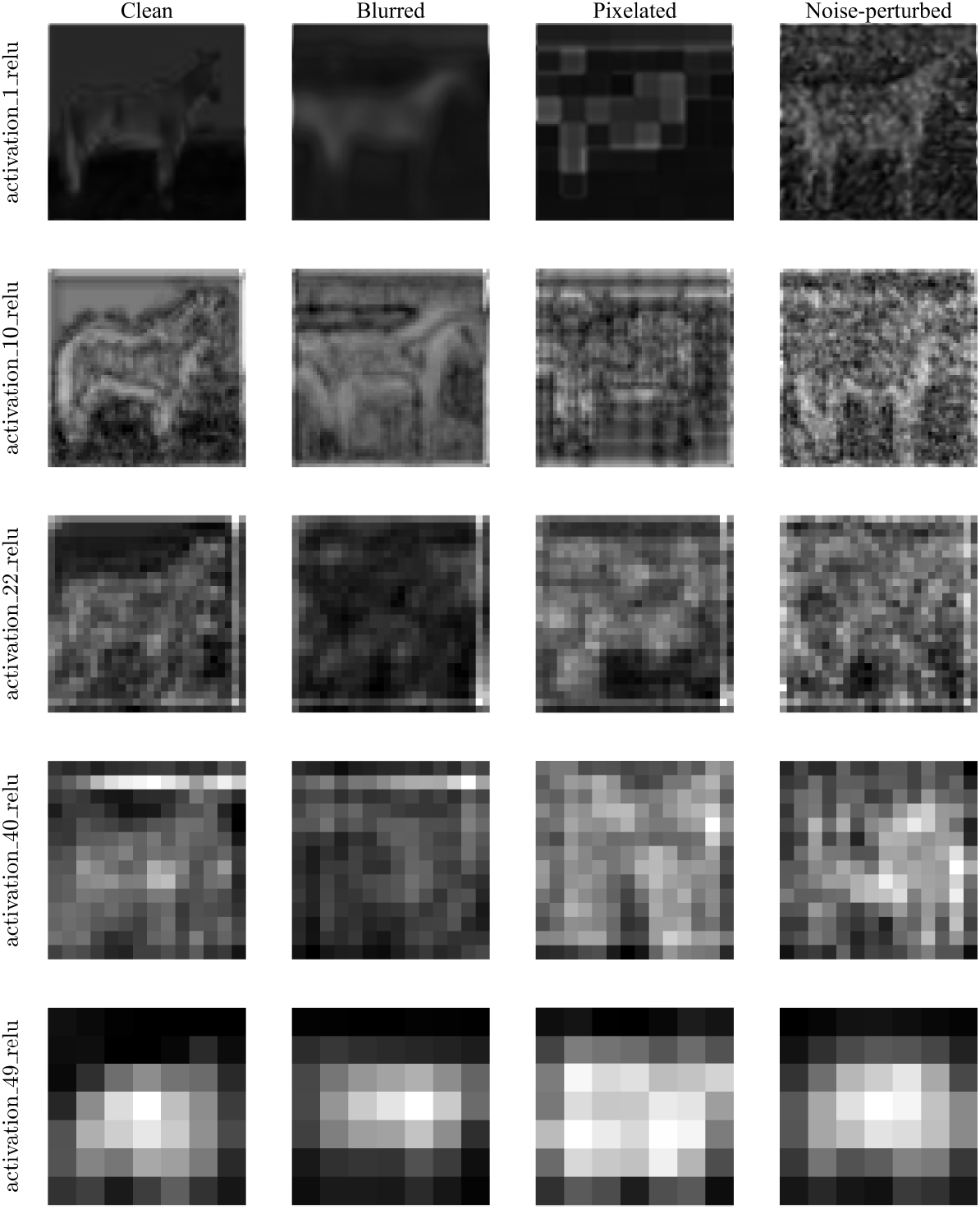
Qualitative example of layer-wise activation structure for one image under different nuisance transformations. Columns show the clean, blurred, pixelated, and noise-perturbed versions of the same input image. Rows show a subset of the analysed ResNet-50 layers: `activation_1_relu, activation_10_relu, activation_22_relu, activation_40_relu, and activation_49_relu`. The figure illustrates how distortion-dependent structure remains visible across the representational hierarchy and complements the information-theoretic summaries in Figure 7.

In the clean-only object run, object-relevant information increased across the network hierarchy (Figure 7A). Object-class information increased from approximately 0.11 in the first analysed layer to approximately 0.59 at `avg_pool`. Superordinate animal–vehicle information increased more strongly, from approximately 0.22 to approximately 0.82. The environment- or affordance-like target also increased, from approximately 0.13 to approximately 0.54. Because no image-condition variation was present in this run, nuisance information was treated as zero by convention. Thus, when the stimulus set contained only clean images, unsupervised activation-space partitions increasingly preserved object-relevant information across the network hierarchy. This pattern is consistent with the general view that later layers in deep visual networks tend to support more abstract object-relevant representations [35, 32]. In the pooled nuisance run, the pattern changed qualitatively (Figure 7B). The image set contained clean, blurred, pixelated, and noise-perturbed versions of the CIFAR-10 stimuli. Under these conditions, nuisance condition became the dominant preserved target after the first analysed layer. Nuisance information rose to approximately 0.91 at `activation_10_relu`, reached approximately 0.96 at `activation_22_relu`, and remained high through `avg_pool` at approximately 0.89. In contrast, object-class, superordinate animal–vehicle, and environment- or affordance-like information remained much lower throughout the same run. The layer-derived category map was therefore highly informative, but primarily about the nuisance variable rather than the semantic object targets. This result is compatible with the broader concern that activation-space structure can be strongly shaped by image statistics, transformations, or nuisance variation rather than by semantic object category alone [31, 38]. The final-layer comparison makes the contrast between scenarios explicit (Figure 7C). At `avg_pool`, the clean-only run preserved substantial object-relevant information, especially for the superordinate animal–vehicle target. In the pooled nuisance run, object-relevant information was much weaker, whereas nuisance condition was strongly preserved. The same network, same clustering logic, and same information-theoretic evaluation therefore produced different category-map relevance depending on the structure of the stimulus set. When nuisance transformations were absent, object-relevant structure was recovered. When nuisance transformations were pooled with object variation, the unsupervised category map preferentially captured transformation-induced nuisance structure.

This neural-network extension provides an important qualification to the synthetic demonstrations. ResNet-50 clearly contains object-relevant information, as shown by the clean-only run. However, when category maps are derived by unsupervised clustering of activation geometry in a stimulus set with strong nuisance transformations, the dominant preserved variable need not be semantic object class. The analysis therefore separates three issues that are often conflated: a representation can be high-dimensional or structured, a category map can have substantial entropy, and the information preserved by that map can still be irrelevant to the intended semantic target. This distinction is central to information-theoretic approaches in which compression is evaluated by the information retained about a specified target variable, not by compression or separability alone [22, 23].

## 4 Discussion

The present analysis supports a simple but important conclusion: categorisation cannot be evaluated by compression alone. A category map reduces the complexity of a stimulus space, but its usefulness depends on what information is preserved by that reduction. Category entropy measures how much categorical structure remains after compression, whereas mutual information with a target variable measures whether that remaining structure is relevant to a specified cognitive, behavioural, or computational target [19, 20, 22, 23]. The synthetic demonstrations showed that the same stimulus space can support multiple candidate category maps with sharply different informational consequences. Identity-preserving, action-preserving, geometric, nuisance-based, random, and one-category maps all impose partitions on the same points. Some preserve the target of interest; others do not. The difference cannot be read off from category entropy alone. A random partition can have high entropy and low usefulness. A one-category partition can achieve maximal compression and no usefulness. A nuisance partition can be highly informative, but only about a nuisance target. This result is consistent with earlier accounts in which category utility depends on the relation between category structure and informative properties, rather than on category structure alone [3, 4]. This distinction is directly relevant to theories of category learning. Classical work on basic-level categories emphasised that some levels of abstraction are especially useful because they balance informativeness and economy [6, 7, 2]. The present formulation expresses this idea in information-theoretic terms. A category level is useful when it preserves the target variable required by the task while avoiding unnecessary category complexity. Subordinate, basic, and action/superordinate-like levels therefore need not be treated as fixed cognitive levels with universal value. They can be treated as alternative compressions whose value depends on the target structure that must be retained. This also aligns with computational and rational accounts in which categories support generalisation by encoding assumptions about which properties should transfer across instances [9, 10, 5, 12].

The neural-network extension strengthens this conclusion in a learned representational setting. In the clean-only ResNet-50 run, layer-derived category maps increasingly preserved object-relevant information across the network hierarchy. In the pooled nuisance run, however, the same analysis preferentially recovered nuisance condition. The contrast is important because it shows that the framework does not merely declare neural-network clusters to be informative or uninformative. Rather, it identifies which target variable is preserved by a given partition of activation space. This is relevant to broader work using deep networks as models of visual representation and as comparison systems for biological object recognition [33, 36, 35, 32]. The result also clarifies the role of target specification. If the relevant target is nuisance condition, the deeper ResNet-50 category maps in the pooled run are highly informative. If the relevant target is object class, superordinate animal–vehicle class, or an affordance-like grouping, the same maps are much less informative. Conversely, when nuisance variation is removed in the clean-only run, final-layer maps preserve substantially more object-relevant information. Therefore, the usefulness of a categorisation cannot be inferred from entropy or clustering structure alone. It depends on the relation between the category map and the variable that the system is required to preserve. The neural-network result should not be interpreted as evidence that ResNet-50 lacks object information. The clean-only analysis shows the opposite. Rather, the pooled nuisance result shows that an unsupervised clustering procedure applied to transformed images can preferentially recover the dominant variance structure of the activation space. Under the present stimulus construction, this dominant structure is nuisance-related. This interpretation is consistent with concerns that neural-network representations can be strongly shaped by image statistics, texture, or transformation structure, rather than by semantic object category alone [31, 38]. A supervised readout, conditional mutual information analysis, or object-information analysis within each nuisance condition would be required to determine how much object-relevant information remains available once transformation-induced variance is controlled.

Several limitations follow from the deliberately minimal design. The synthetic demonstrations use low-dimensional Gaussian stimulus spaces and explicitly defined target variables. This is useful for isolating the formal logic, but it does not capture the full structure of natural categories. Real categories involve multi-feature distributions, hierarchical dependencies, contextual variation, causal knowledge, embodiment, and task-specific constraints [42, 2]. The present analysis should therefore be understood as a formal demonstration of a principle rather than a complete model of natural category learning. The neural-network extension is also exploratory. CIFAR-10 images are small, and the nuisance transformations are visually strong [40]. Moreover, unsupervised clustering of activation vectors is only one way to derive category maps from a network. Other analyses could use supervised probes, conditional mutual information, representational similarity analysis, invariance metrics, or downstream behavioural readouts [34, 35]. Nevertheless, the result is informative because it reveals exactly the problem addressed by the manuscript: structured compression can preserve a variable that is not aligned with the intended target.

Future work can extend the framework in several directions. First, target-conditional information measures could be used to ask whether object information is preserved within each nuisance condition:

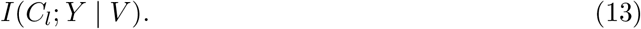

Second, category maps could be compared across different levels of a network, different architectures, or different training objectives. Third, behavioural and neural data could be analysed using the same framework by treating observed responses, neural population states, or model activations as candidate category maps. This may be particularly useful for comparative cognition, where the relevant question is often not whether an animal possesses a rich human-like concept, but whether a nervous system preserves enough target-relevant information to support a specific discrimination. In biological terms, this can be framed as a form of functional pleiotropy or exaptive reuse: a representational system shaped by one ethologically important demand may also preserve information useful for another, thereby supporting discrimination along a target dimension that was not necessarily the original basis of selection [43]. The present framework could formalise such cases by asking how much information about a novel target is preserved by a category map that may have originally been shaped by a different behavioural demand. Fourth, category learning could be modelled as the search for a category map that maximises target preservation under a complexity constraint. Such an extension would connect the present analysis more directly to information-bottleneck approaches and computational models of category learning [22, 23, 11]. The central implication is that category maps should be understood as target-preserving compressions. Compression is necessary for categorisation, but it is not sufficient. A useful category system preserves the information that matters for the organism, task, or model under consideration. Entropy describes how much category structure remains. Mutual information with a specified target describes whether that structure is useful.

## 5 Conclusions

Categorisation should not be evaluated by compression alone. A category map may reduce stimulus complexity while preserving little information about the variable that matters for recognition, prediction, or control. The present analyses showed this principle in both synthetic stimulus spaces and learned visual representations. Across the synthetic demonstrations, the same stimulus space supported multiple category maps with different informational consequences, and their usefulness depended on which target variable was to be preserved. In the neural-network extension, unsupervised layer-derived category maps preserved object-relevant information in a clean-only setting, but preferentially preserved nuisance condition when strong transformation variation was pooled into the stimulus set. These findings support an information-theoretic view of categorisation as target-preserving compression: the value of a category system depends not only on how strongly it compresses a stimulus space, but on what information remains available after that compression.

## 6 Author contribution

C.D.D. conceived the study, developed the methodology, implemented the numerical analyses, generated the figures, and wrote and revised the manuscript.

## 8 Institutional review

This work is purely computational and does not involve human participants, animals, or identifiable personal data.

## 9 Data availability

All data analysed in this study were either generated synthetically within the modelling pipeline or derived from the publicly available CIFAR-10 dataset processed through a pretrained ResNet-50 network. The MATLAB code used to construct the synthetic demonstrations, extract and analyse the layer-derived category maps, compute the information-theoretic summaries, and generate all figures is openly available at https://github.com/ChristophDahl/entropic-compression-category-maps under the MIT License. The repository includes a master script that reproduces the analyses and figures reported in this manuscript.

